# Impact of an Urban Sanitation Intervention on Enteric Pathogen Detection in Soils

**DOI:** 10.1101/2021.04.02.438233

**Authors:** Drew Capone, David Berendes, Oliver Cumming, David Holcomb, Jackie Knee, Konstantinos T. Konstantinidis, Karen Levy, Rassul Nalá, Benjamin B. Risk, Jill Stewart, Joe Brown

**Author notes:** Corresponding author: Joe Brown, Address: 135 Dauer Dr, Chapel Hill, NC 27599, USA.

## Abstract

Environmental fecal contamination is common in many low-income cities, contributing to a high burden of enteric infections and associated negative sequelae. To evaluate the impact of a shared onsite sanitation intervention in Maputo, Mozambique on enteric pathogens in the domestic environment, we collected 179 soil samples at shared latrine entrances from intervention (n= 49) and control (n= 51) compounds during baseline (pre-intervention) and after 24 months (post-intervention) as part of the Maputo Sanitation Trial. We tested soils for the presence of nucleic acids associated with 20 enteric pathogens using a multiplex reverse transcription qPCR platform. We detected at least one pathogen-associated target in 91% (163/179) of soils and a median of 3 (IQR=1.5, 5) pathogens. Using a difference-in-difference analysis and adjusting for compound population, visibly wet soil, sun exposure, wealth, temperature, animal presence, and visible feces, we estimate the intervention reduced the probability of ≥1 pathogen detected by 15% (adjusted prevalence ratio, aPR=0.85; 95% CI: 0.70, 1.0) and the total number of pathogens detected by 35% (aPR =0.65; 0.44, 0.95) in soil 24 months following the intervention. These results suggest that the intervention reduced the presence of some fecal contamination in the domestic environment, but pathogen detection remained prevalent 24-months following the introduction of new latrines.

## INTRODUCTION

Onsite sanitation systems are designed to sequester human feces away from human contact and prevent the transport of fecal-oral pathogens through well-understood transmission pathways.^1^ Large-scale, rigorous randomized controlled trials (RCTs) of onsite sanitation systems – including sanitation alone and combinations of water, sanitation, and hygiene (WASH) interventions – have found mixed effects on health outcomes, such as diarrhea and child growth.^2–7^ Assessing the impact of WASH interventions on enteric pathogens in the environment can improve our understanding of pathogen transmission from an infected individual to a new host via the environment, a core intermediate outcome of these trials. Such data may help explain why some WASH interventions observed improved health outcomes and others did not.^8^

There is a growing body of literature that soils contaminated by feces in public and domestic environments pose infection risks.^9–13^ In health impact trials that assess improved onsite sanitation systems, soils are assessed to measure how effectively the intervention sequestered human feces.^14–18^ Latrines and septic tanks are useful barriers against the transport of human feces into the environment. However, enteric pathogens may still move into soils through open defecation^19^, unhygienic pit emptying^20,21^, fecally contaminated greywater^22,23^, improper disposal of children’s feces or anal cleansing materials^24,25^, latrine flooding^20,26,27^, animal feces^28–30^, or subsurface transport from unlined pits^31–33^. Domestic soils contaminated by enteric pathogens can pose infection risks beyond incidental^34^ and direct^35^ soil ingestion: contaminated soil may be transported to hands, food, fomites, or household stored water.^36^ For these reasons, soils may be a useful matrix to assess the impact of onsite sanitation interventions.

Detecting enteric pathogens via molecular methods is increasingly used to assess the impact of WASH interventions on the transport of these pathogens through the environment.^37–39^ Molecular detection of pathogens offers additional insights, as health impact studies have historically relied on fecal indicator bacteria (FIB), as a proxy for enteric pathogens for reasons of cost, capacity and feasibility.^17,36, 40–42^ However, a 2016 meta-analysis^43^ found that improved sanitation had no effect on the presence of FIB in the environment, possibly because these indicators are often pervasive in low-income settings^15,16,36, 44–46^ and common FIB, like *E. coli*, may be naturalized in the environment^47–49^.

The Maputo Sanitation (MapSan) Trial was the first rigorous controlled before-and-after trial to evaluate the effect of an urban onsite sanitation intervention on child health.^24,50,51^ We conducted the trial in low-income, informal neighborhoods in Maputo, Mozambique, where WASH conditions are poor, and the burden of enteric disease is high.^20,24,44,52^ Water and Sanitation for the Urban Poor (WSUP, a non-governmental organization) delivered the intervention to compounds composed of household clusters that shared sanitation and courtyard space. The intervention was built inside the compound boundary and was part of the households’ living environment. WSUP replaced shared onsite sanitation systems in poor condition with pour-flush toilets that included septic tanks and soak-away pits (Text S1). Control compounds were concurrently enrolled from the same or adjacent neighborhoods as intervention compounds and continued using existing shared sanitation infrastructure. Detailed descriptions of the inclusion criteria for intervention and control compounds are described elsewhere.^20,24^

A latrine entrance is an ideal soil sampling location to determine the effectiveness of onsite sanitation interventions because it is a standardized location near the fecal waste in the containment chamber.^15,16,53^ Soils in low-income Maputo are characterized as coarse to fine sand or silty sand.^54^ While the fate and transport of pathogens through soils is dependent on the individual pathogen and environmental conditions^55^, the high porosity of Maputo’s sandy soils combined with a high water table in the study area^44^ offers potential for pathogen movement.^56^ This high risk of fecal contamination suggests we could plausibly observe a reduction in enteric pathogens in soil at latrine entrances if the intervention infrastructure performed better than controls at safely containing fecal wastes.^57^ Our study aim was to assess if the intervention reduced the detection of ≥1 pathogen, the total number of pathogens, or any individual pathogen in latrine entrance soils from MapSan intervention compounds compared to controls.

## MATERIALS AND METHODS

### Sample Collection

We prospectively collected latrine entrance soil samples – defined as a location one-meter away from the latrine entrance in the direction of entry or the nearest point not covered by cement – from 49 intervention and 51 control compounds at baseline (pre-intervention) and from the same compounds 24-months following the intervention, for a total of 200 samples (Text S2). We defined this sample location *a priori* as one that could be standardized across all compounds in the study. Using a spade and ruler, we scooped a 10 cm x 10 cm x 1 cm volume of soil into a Whirl-Pak^®^ bag (Nasco, Fort Atkinson, WI). The spade and ruler were sterilized between uses with 10% bleach and 70% ethanol. At the time of sampling, enumerators recorded whether the soil was visibly wet and estimated the daily sun exposure (full sun, partially shaded, full shade).^44^ Samples were stored on ice for transport to the Ministry of Health in Maputo, Mozambique, frozen at −20°C for approximately six months, aliquoted into 2 ml cryovials while working on dry ice, and then stored at −80°C. During storage at −20°C, some samples (n = 21) were unable to be evaluated because the permanent marker labeling on some Whirl-Pak^®^ bags wore off and some bags burst open. All aliquoted samples (n = 179) were shipped from the Mozambican Ministry of Health in Maputo, Mozambique to Atlanta, GA, USA on dry ice (−80° C) with temperature monitoring for molecular analysis. We obtained compound observation data and socioeconomic characteristics from the MapSan baseline and 24-month survey datasets, which were collected concurrent to soil samples.^24,58^

### Sample Processing

At Georgia Institute of Technology in Atlanta, GA, USA, we incubated 250 mg of each soil sample at 105°C for 1 hour to determine moisture content^13,59^, then discarded the dry soil. We then extracted total nucleic acids from a separate 1-gram (calculated for dry weight) portion of each sample, and spiked samples with MS2 (Luminex Corporation, Austin, TX) as an extraction control. Following the manufacturer’s protocol, we extracted RNA using the RNeasy PowerSoil Total RNA Kit and DNA using the RNeasy PowerSoil DNA Elution Kit (Qiagen, Hilden, Germany). On each day of extraction (approximately every 5-15 samples), we included one negative extraction control (sterile deionized water). We tested sample extracts for matrix inhibition using the Applied Biosystems Exogenous Internal Positive Control Assay^60^ (Applied Biosystems, Waltham, Massachusetts) before downstream molecular analysis (Text S3).

We assayed extracted nucleic acids from all samples using a custom TaqMan Array Card (TAC) (ThermoFisher Scientific, Waltham, MA) that tested for 20 enteric pathogens in duplicate wells following Liu *et al*. 2013^61^, including ten bacteria (*Campylobacter jejuni/coli*, *Clostridium difficile* [*tcdA* and *tcdB* gene], Enteroaggregative *E. coli* [EAEC, *aaiC* and *aatA* gene], *Shigella/*Enteroinvasive *E. coli* [EIEC, *ipaH* gene], Enteropathogenic *E. coli* [EPEC, *bfpA* and *eae* gene], Enterotoxigenic *E. coli* [ETEC, heat-labile and heat-stabile enterotoxin gene], Shiga-toxin producing *E. coli* [STEC, *stx1* and *stx2*], *Salmonella* spp., *Vibrio cholerae*, and *Yersinia* spp.), five viruses (adenovirus 40/41, astrovirus, norovirus [GI and GII], rotavirus A, and sapovirus [I, II, IV, and V],), three protozoa (*Cryptosporidium parvum*, *Entamoeba histolytica*, and *Giardia duodenalis*) and two soil-transmitted helminths (*Ascaris lumbricoides*, *Trichuris trichiura*) (Text S4, Table S1, Table S2).^62^ We combined and then added 25 µL of RNA eluant, 25 µL of DNA eluant, and 50 µL of mastermix into each TAC port. We included a positive and negative control on each TAC. The positive control was a plasmid that included all assay gene sequences and the negative control was either extract from a negative extraction control or sterile water.^63^ The thermocycling conditions were as follows: 45°C for 10 minutes and 94°C for 10 minutes, followed by 45 cycles of 94°C for 30 seconds and 60°C for 1 minute, with a ramp rate of 1°C/second between each step. We visually compared exponential curves and multicomponent plots with the positive control plots to validate positive amplification^12^; positive amplification in one or both duplicate wells below a quantification cycle (Cq) of 40 was called as a positive for a target (Text S4).^62,64^

### Data analysis

We analyzed data in R version 4.0.0 (R Foundation for Statistical Computing, Vienna, Austria). We used a difference-in-difference (DID)^65^ approach to assess the impact of the intervention – our exposure variable – on our outcomes compared to the control group. Our outcomes included the detection (i.e., binary presence/absence) of ≥1 of the enteric pathogens measured, the total number of pathogens detected out of 20, and a separate analysis for each pathogen individually. We used generalized estimating equations (GEE)^66^ to fit unadjusted and adjusted Poisson regression models with robust standard errors, with an exchangeable correlation structure. We accounted for clustering between compounds across the two study phases because the intervention was implemented at the compound level.^67^

To generate adjusted estimates, we selected nine covariates from the MapSan baseline and 24-month datasets based on their biological plausibility to impact the transport^57^ or persistence^68^ of pathogens in the domestic environment and previously reported associations in the literature^36,44^ (Table S3). We used the same nine covariates to adjust all DID models: compound population (a 10-person increase in compound population), wealth (one-quartile increase in wealth index^69^), soil moisture (assessed visually at the time of sampling), sun exposure status (estimated at the time of sampling; full sun, partially shaded, shaded^44^), the mean-centered average air temperature in Fahrenheit for the day of and day preceding sample collection (i.e., two-day average), a binary variable for the presence of cats, a binary variable for the presence of dogs, a binary variable for the presence of chickens or ducks, and a binary variable for the presence of visible animal or human feces in the compound (Table S4).

To estimate the intervention’s effect, we used the interaction of dummy variables representing treatment status (intervention vs. control) and trial phase (baseline or 24-month). Consequently, we present the effect estimates from our DID analysis as ratio measures (ratio of prevalence ratios, PR) instead of absolute differences. We fit separate GEE models to measure the association between intervention status and the detection of ≥1 pathogen and the total number of pathogens detected among the 20 targets we identified *a priori*. Likewise, we fit DID models to estimate the intervention’s impact for each individual pathogen assessed, but we excluded any pathogen not detected in at least 5% of control and intervention samples during both phases.

### Ethics

The study protocol was approved by the Comité Nacional de Bioética para a Saúde (CNBS), Ministério da Saúde (333/CNBS/14), the Research Ethics Committee of the London School of Hygiene and Tropical Medicine (reference # 8345), and the Institutional Review Board of the Georgia Institute of Technology (protocol # H15160). The overall trial was pre-registered at ClinicalTrials.gov (NCT02362932), but we did not pre-register this environmental analysis.

## RESULTS

### Matched samples

We analyzed latrine entrance soils collected at baseline from 48 control compounds and 43 intervention compounds, and soils collected at the 24-month phase from 45 control and 43 intervention compounds (Table S4). We did not analyze twelve intervention samples and nine control samples because they were either lost or damaged during storage. This resulted in some samples collected at either phase not having a matched sample from the same compound from the earlier or later phase. Among the 93 control samples analyzed, 42 compounds had samples from both phases (n=84), six baseline samples did not have a matched 24-month phase sample, and three 24-month samples did not have a matched baseline sample. Among the 86 intervention samples analyzed, 41 compounds had samples from both phases (n=82), two baseline samples did not have a matched 24-month phase sample, and two 24-month samples did not have a matched baseline sample. There was a mean of 788 days between the collection of matched control samples (sd = 36, min = 733, max = 860) and a mean of 789 days between matched intervention samples (sd = 56, min = 731, max = 953). Control and intervention samples were collected approximately during the same period of the year (Figure S1).

### Compound characteristics

Control and intervention compounds had similar wealth indices at baseline (mean= 0.47 [sd=0.09] and mean=0.46 [sd=0.09], respectively, p=0.49) but control compounds had higher wealth indices at the 24-month phase (mean=0.46 [sd=0.12] and mean=0.40 [sd=0.09], respectively, p=0.05) (Table 1). The number of residents in the intervention compounds was greater than control compounds at baseline (mean=19 [sd=7.8] and mean=14 [sd=6.4], respectively, p=0.004) and at the 24-month phase (mean=16 [sd=7.9] and mean=13 [sd=7.0], respectively, p=0.02) (Table 1).

**Table 1.** Characteristics of MapSan trial compounds and households selected for soil sampling.

|  |  |  | Baseline |  |  |  | 24-Month Phase |  |  |  |
| --- | --- | --- | --- | --- | --- | --- | --- | --- | --- | --- |
|  |  |  | Control |  | Intervention |  | Control |  | Intervention |  |
| Characteristic | Level | Metric | N | Summary | N | Summary | N | Summary | N | Summary |
| Wealth index (0-1) | household | mean (sd) | 48 | 0.47 (0.09) | 43 | 0.46 (0.09) | 45 | 0.44 (0.12) | 43 | 0.40 (0.09) |
| Compound population | compound | mean (sd) | 48 | 14 (6.4) | 43 | 19 (7.8) | 45 | 13 (7.0) | 43 | 16 (7.9) |
| Any animal(s) present | compound | n (%) | 48 | 28 (58%) | 43 | 28 (65%) | 45 | 32 (71%) | 43 | 35 (81%) |
| Cat(s) present | compound | n (%) | 48 | 24 (50%) | 43 | 23 (53%) | 45 | 32 (71%) | 43 | 30 (70%) |
| Chicken(s) or duck(s) present | compound | n (%) | 48 | 6 (13%) | 43 | 7 (16%) | 45 | 4 (8.9%) | 43 | 8 (19%) |
| Dog(s) present | compound | n (%) | 48 | 3 (6.3%) | 43 | 4 (9.3%) | 45 | 9 (20%) | 43 | 10 (23%) |
| Other animal(s) present | compound | n (%) | 48 | 1 (2.1%) | 43 | 2 (4.7%) | 45 | 1 (2.2%) | 43 | 0 (0%) |
| Visible feces | compound | n (%) | 48 | 22 (46%) | 43 | 22 (51%) | 45 | 4 (8.9%) | 43 | 4 (9.3%) |
| Visibly wet soil | sample | n (%) | 48 | 37 (77%) | 43 | 34 (79%) | 45 | 37 (82%) | 43 | 34 (79%) |
| Partially shaded soil | sample | n (%) | 48 | 24 (50%) | 43 | 13 (30%) | 45 | 30 (67%) | 43 | 28 (65%) |
| Fully shaded soil | sample | n (%) | 48 | 14 (29%) | 43 | 20 (47%) | 45 | 10 (22%) | 43 | 9 (21%) |
| Temperature (°F) | date | mean (sd) | 48 | 72 (4.5) | 43 | 70 (4.3) | 45 | 72 (4.7) | 43 | 73 (5.3) |
| No useable sanitation infrastructure | compound | n (%) | 48 | 3 (6.3%) | 43 | 4 (9.3%) | 45 | 0 (0%) | 43 | 0 (0%) |
| Pit latrine with slab | compound | n (%) | 48 | 27 (56%) | 43 | 14 (14%) | 45 | 18 (40%) | 43 | 0 (0%) |
| Pit latrine without slab | compound | n (%) | 48 | 16 (33%) | 43 | 24 (56%) | 45 | 14 (31%) | 43 | 0 (0%) |
| Pour-flush toilet (non-intervention) | compound | n (%) | 48 | 2 (4.2%) | 43 | 1 (2.2%) | 45 | 13 (29%) | 43 | 0 (0%) |
| Intervention infrastructure | compound | n (%) | 48 | 0 (0%) | 43 | 0 (0%) | 45 | 0 (0%) | 43 | 43 (100%) |

Note: Wealth index created using the 2013 Simple Poverty Scorecard© for Mozambique

Reported or observed animal ownership was high across trial arms during both phases (Table 1). Most compounds had at least one animal at baseline (62% [56/91]) including cats (50% [24/48] control, 53% [23/43] intervention), chickens or ducks (13% [6/48] control, 16% [7/43] intervention), and dogs (6.3% [3/48] control, 9.3% [4/43] intervention). Three-quarters of compounds had at least one animal 24-months post intervention (76% [67/88]): cats were most common (71% [32/45] control, 70% intervention [30/43]), followed by dogs (20% [9/45] control, 23% [10/43] intervention), and chickens or ducks (8.9% [4/45] control, 19% [8/43] intervention).

At baseline seven compounds had no useable sanitation infrastructure (6.3% [3/48] control, 9.3% [4/43] intervention) and three compounds had pour-flush sanitation (4.2% [2/48] control, 2.3% [1/43] intervention) (Table 1). Control compounds more often had pit latrines with slabs (56%, [27/48]) than without slabs (33%, [16/48]), compared to intervention compounds, which more often had pit latrines without slabs (56%, [24/43]) than with slabs (33%, [14/43]) (p=0.09). At the 24-month phase, most control compounds had a pit latrine (with slab 40%, [18/45]; without slab 31%, [14/45]), but some (29%, [13/45]) had independently upgraded their pit latrines to pour-flush toilets. All intervention compounds (100%, [43/43]) still had the intervention sanitation infrastructure at the 24-month phase.

### Laboratory Controls

We did not observe inhibition in any sample (Text S3). We observed positive amplification for all assays using our positive controls (n = 32). We did not observe positive amplification for any assay in our extraction controls (n=16), nor any template controls (n=16) below a Cq of 40.

### All Pathogens

We detected at least one pathogen in 91% (163/179) of latrine entrance soils, two or more pathogens in 75% (134/179), and a mean of 3.4 out of 20 measured targets (IQR=3.5). The four most frequently detected pathogens were *Ascaris lumbricoides* (62%, [111/179]), EAEC (46%, [82/179]), *Giardia duodenalis* (36%, [64/179]), and astrovirus (26%, [47/179]). We found evidence that the intervention reduced the detection of ≥1 pathogen in latrine entrance soils by 15% (aPR = 0.85, 95% CI [0.70, 1.0]) and the total number of pathogens by 35% (aPR = 0.65, 95% CI [0.44, 0.95]) (Table 2). The mean Cq values of detected pathogens were similar across trial arms and phases (Table S5).

**Table 2.**
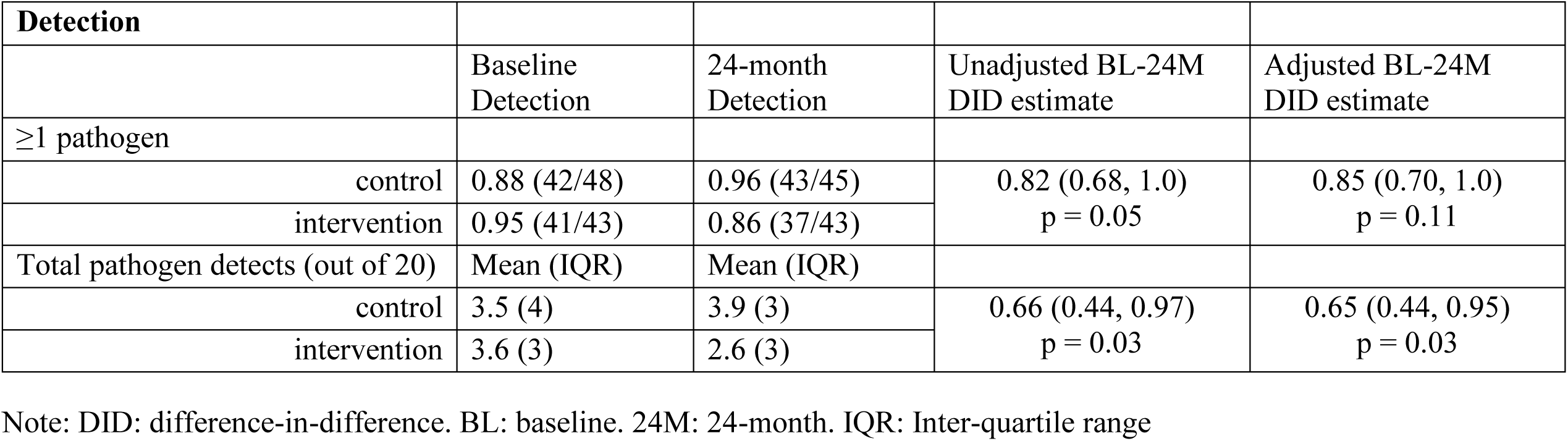
Detection of pathogens at baseline and 24-month.

| Detection |  |  |  |  |
| --- | --- | --- | --- | --- |
|  | Baseline<br>Detection | 24-month<br>Detection | Unadjusted BL-24M<br>DID estimate | Adjusted BL-24M<br>DID estimate |
| ≥1 pathogen |  |  |  |  |
| control | 0.88 (42/48) | 0.96 (43/45) | 0.82 (0.68, 1.0)<br>p = 0.05 | 0.85 (0.70, 1.0)<br>p = 0.11 |
| intervention | 0.95 (41/43) | 0.86 (37/43) |  |  |
| Total pathogen detects (out of 20) | Mean (IQR) | Mean (IQR) |  |  |
| control | 3.5 (4) | 3.9 (3) | 0.66 (0.44, 0.97)<br>p = 0.03 | 0.65 (0.44, 0.95)<br>p = 0.03 |
| intervention | 3.6 (3) | 2.6 (3) |  |  |

Note: DID: difference-in-difference. BL: baseline. 24M: 24-month. IQR: Inter-quartile range

There was a consistent trend among all individual pathogens except for astrovirus: the adjusted point estimates for nine of the ten most frequently detected had point estimates below 1.0 (Table 3). The confidence intervals around the adjusted DID estimates of effect were also below 1.0 for three pathogen targets: *Ascaris lumbricoides* (aPR = 0.62, 95% CI [0.39, 0.98]), EAEC (aPR=0.51, 95% CI [0.27, 0.94]), and EPEC (aPR = 0.20 95% CI [0.05, 0.82]).

**Table 3.**
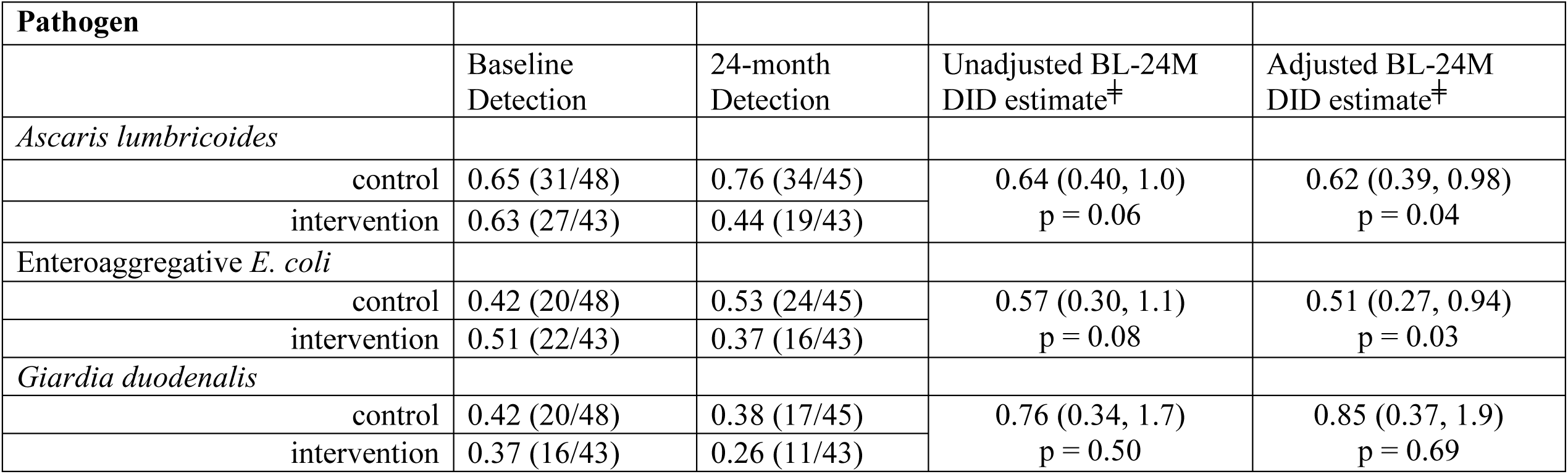
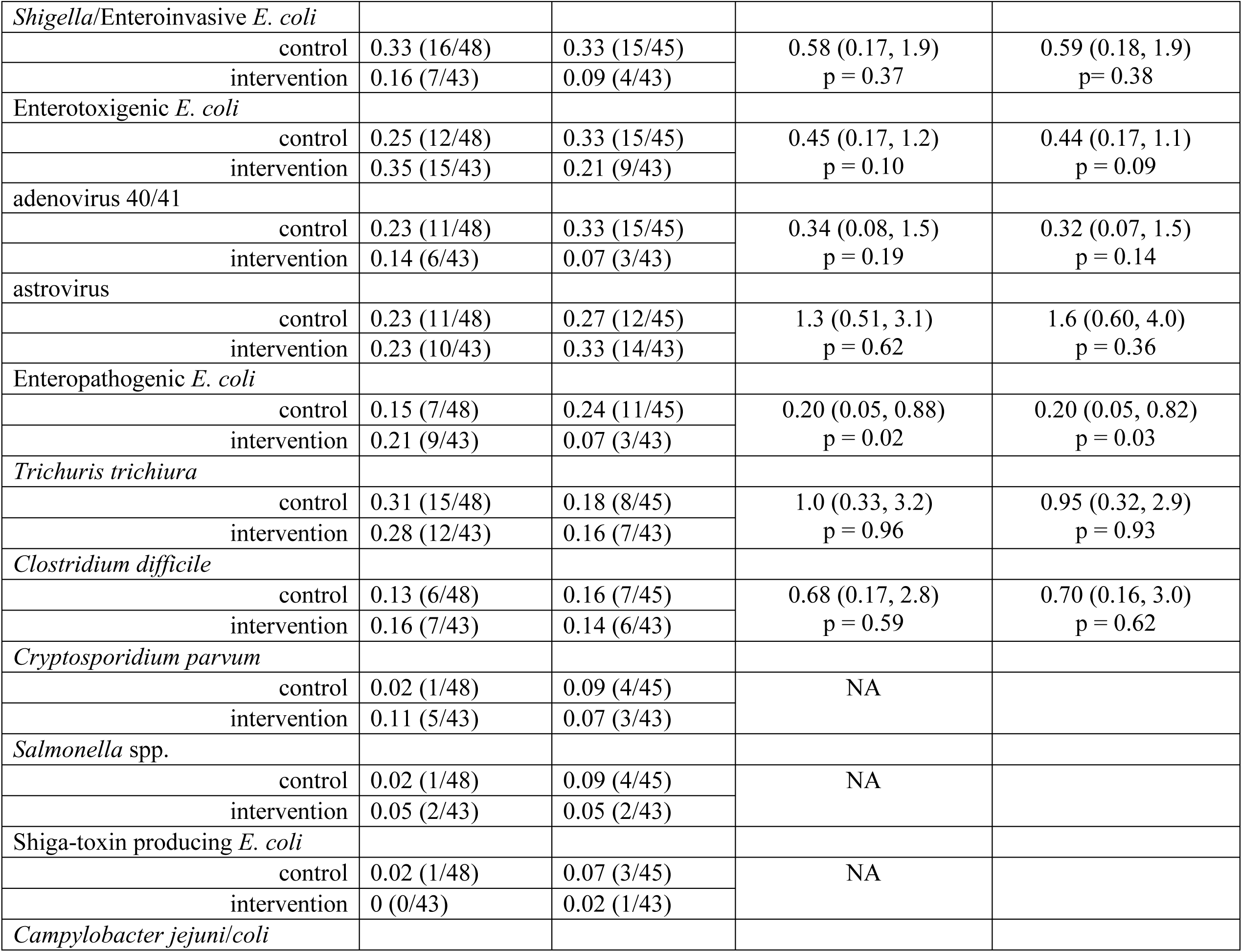
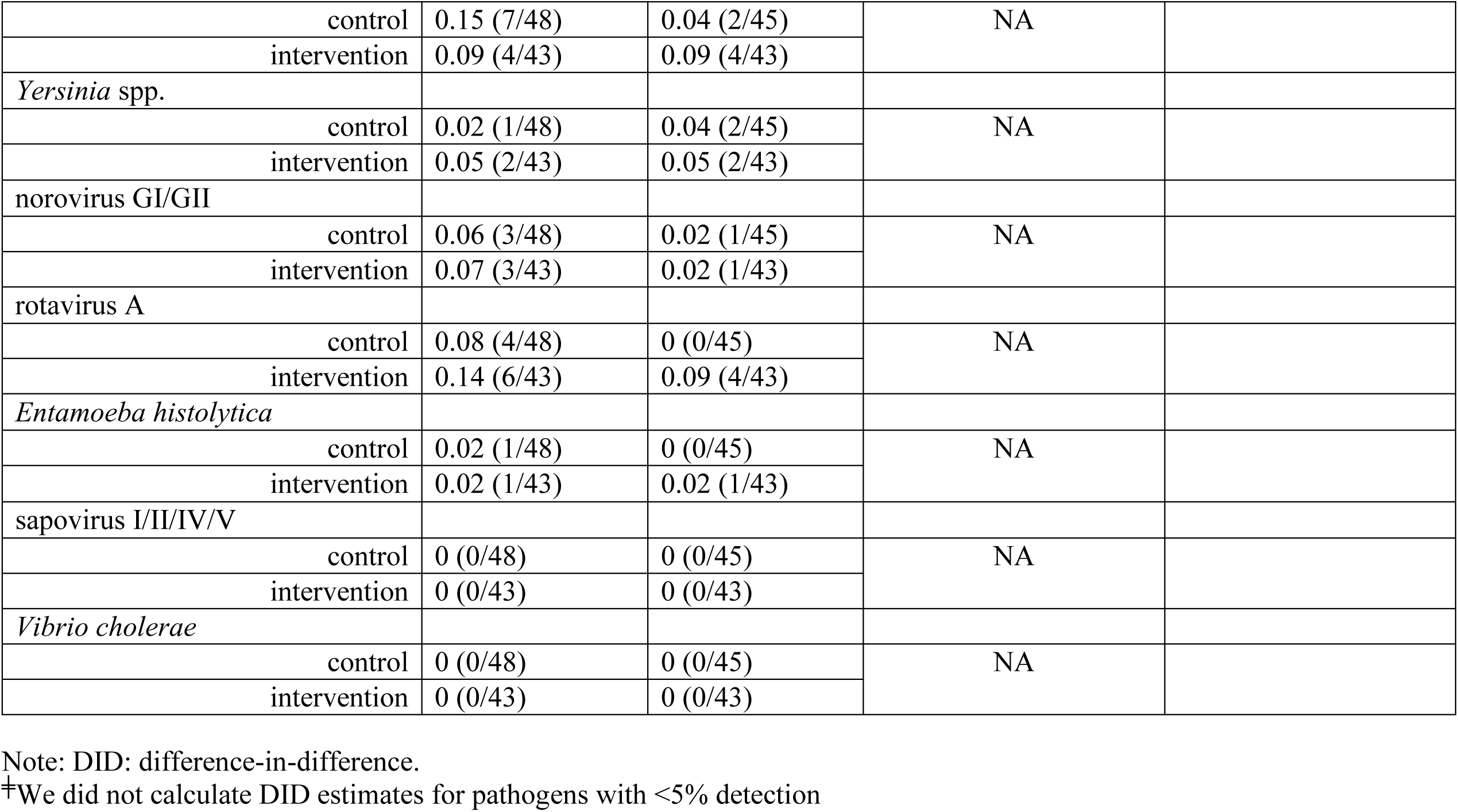
Detection of individual pathogens at baseline and 24-month. Sorted by detection in control soils at the 24-month phase.

## DISCUSSION

We found evidence that the onsite shared urban sanitation intervention evaluated in the MapSan trial was somewhat protective against the detection of ≥1 pathogen and against the total number of pathogens in latrine entrance soils. The adjusted estimates for nine of the ten most common pathogens were consistently protective (DID estimates = 0.20-0.95) and pathogen-specific effect estimates from adjusted models were protective for *Ascaris lumbricoides*, EAEC, and EPEC. This suggests that intervention septic tanks may have better sequestered or inactivated these pathogens, which are passed in stool, compared with controls.

Most of the other pathogens we frequently detected in soils were measured in child stools via multiplex end-point PCR as part of the MapSan trial, with the exception of EAEC, EPEC, and astrovirus. At baseline, *Shigella*/EIEC (44%) and *Trichuris trichiura* (37%) – generally thought to be transmitted human-to-human – were the second and third most common pathogens detected in child stool^24,50^, following *Giardia* (51%) which can be zoonotic^70^. Given the high prevalence of anthroponotic enteric pathogens in stools and the lack of a zoonotic reservoir for *Shigella*/EIEC and *Trichuris trichiura*^71,72^, the trial may have had greater power to observe an effect on *Shigella*/EIEC and *Trichuris trichiura* compared with other pathogens. For children born into study compounds before the 24-month visit, the intervention reduced the detection of *Shigella*/EIEC in children’s stools by 51% and *Trichuris trichiura* by 76%.^58^ Results from soils in this study differ from trial findings in stools: while we observed a 41% reduction in *Shigella*/EIEC detection, we identified no difference with respect to detection of *Trichuris trichiura*. This absence of impact on *Trichuris trichiura* in soils may have been due to limited power from infrequent detection; we did observe a reduction in the other STH assessed, *Ascaris lumbricoides*, which was the most frequently detected individual pathogen in soils. The MapSan trial found the sanitation intervention reduced the detection of *Ascaris lumbricoides* by 32% among children born into study compounds before the 24-month visit, but the confidence interval included the null.^58^ Overall, the protective trend we observed in soils, therefore, is consistent with the enteric infection data for children born into trial compounds. This may suggest that the intervention reduced the transport of pathogens to latrine entrance soils, and subsequently contributed to a reduction in children’s exposures, but our small sample size and the resulting uncertainty of point estimates suggest results should be interpreted with caution.

Compared to other recent large-scale, rigorous trials of onsite sanitation improvements in rural Bangladesh (pour flush to double-pit latrine)^2^, rural Kenya (single unlined pit latrine with plastic slab and hole-lid)^3^, and rural Zimbabwe (ventilated improved pit latrine)^4^, we evaluated a more sophisticated intervention that included site-specific engineered septic tanks and subsurface discharge of aqueous effluent to a soakaway pit^24,73^, and it is the only recent controlled health impact trial of onsite sanitation to take place in an urban setting. In the early 2000s, Barreto *et al*. observed health benefits from household sewerage connections in urban Brazil in an uncontrolled trial^74,75^. However, the scope, complexity, and cost of that intervention make it an imperfect point of comparison.

The WASH Benefits Trial (WASH-B) evaluated the impact of single and combined water, sanitation, and handwashing intervention arms in rural Bangladesh and Kenya. In Bangladesh, a molecular analysis of household entrance soils, hand rinses, and stored water from the sanitation arm found no significant reductions in enteric pathogens (EAEC, EPEC, STEC, *Shigella*/EIEC, ETEC, norovirus, *Cryptosporidium* spp., *Giardia duodenalis*) or microbial source tracking markers (HumM2, BacCow).^38^ The combined WASH arm and individual water treatment arm observed a reduction in *E. coli* prevalence and concentration in stored drinking water; the individual water treatment and handwashing arms reduced *E. coli* prevalence and concentration in food. WASH-B trial arms in Bangladesh did not observe reductions in *E. coli* in courtyard soil, ambient waters, child hands, or sentinel objects.^76,77^ Likewise, WASH-B Kenya found the individual water treatment arm and combined WASH arm reduced culturable *E. coli* in stored drinking water, but not along other transmission pathways.^18^ The Sanitation, Hygiene, Infant Nutrition Efficacy Project (SHINE) trial in rural Zimbabwe has not yet published the results from a sub-study on environmental fecal contamination. In separate analyses of environmental samples collected during MapSan baseline^15^ and the 24-month phase^13,21,44^ we found widespread fecal contamination in soils and other environmental compartments. At the 12-month MapSan trial phase Holcomb *et al*. 2021 found the intervention reduced *E. coli* gene densities by more than 1-log_10_ in latrine entrance soils, but observed no reduction in culturable *E. coli* or human microbial source tracking markers.^78^ Our study is the first controlled evaluation of an urban onsite sanitation intervention to show a decrease in the detection of enteric pathogens, via molecular methods, in soils from the domestic living environment.

The intervention may have reduced the presence of enteric pathogens in soils compared with controls because the intervention may have better sequestered or treated fecal material than control latrines. In high-income countries, properly designed, constructed, and maintained septic tank systems have been demonstrated to be efficient and economic alternatives to public sewage disposal systems.^79^ Although some pathogen die-off will occur in pit latrines, the primary purpose of pit latrines is to sequester human feces and reduce exposures, and they are not designed to achieve a specific level of pathogen reduction.^80^ Design features of the intervention septic tanks may have resulted in better treatment of fecal wastes than control systems. Intervention septic tanks contained inlet and outlet pipes configured to maximize detention time, baffles to direct incoming waste downward, t-pipes to ensure sequestration of solids and floatable materials, and a sealed containment chamber to promote anaerobic treatment of stored solids and non-settleable materials. In addition, the intervention septic tank systems represented an upgrade to a more permanent sanitation infrastructure. The construction included masonry block walls, a concrete floor, masonry block lined septic tank, masonry block lined soakaway pit, tin roof, and a water seal squat pan.^20,24,53,73^ These features may have acted as a physical barrier that prevented the contamination of soils by enteric pathogens. At the 24-month phase, most control compounds used a pit latrine with or without a slab, and therefore lacked similar physical barriers such as a water seal. In addition, the control compounds that did upgrade to pour flush sanitation may not have used the same rigorous design criteria as intervention septic tanks.^50^

The extent to which bacterial and viral pathogens may be transported from fecal sludges through the surrounding soil depends on pit characteristics including presence of lining^32^ and the hydrological and soil conditions.^31^ Protozoan cysts and helminth ova are unlikely to be transported out of the pit and into surrounding soil because of their relatively large size.^80–82^ Lateral movement of viral pathogens from unlined or partially lined pit latrines to groundwater has been demonstrated at distances up to 50 meters.^31,83^ This movement is often exacerbated by a high water table^31^, which is present in the study neighborhoods.^53^ While we were unable to assess the lining of control latrines, it is unlikely control linings – if present – matched the construction quality of intervention linings.

Pit latrines in low-income Maputo are often covered when full and rebuilt, or the fecal sludge is emptied and buried or dumped nearby.^20^ The intervention included programming to encourage hygienic pit emptying and provided equipment and training to local organizations to offer hygienic emptying services.^73^ During the 24-month phase, intervention compounds emptied their sanitation systems less frequently and were more likely to have their onsite systems emptied hygienically than control compounds.^20^ Less frequent emptying would have beneficial for two reasons. First, longer residence times would likely have resulted in greater pathogen die-off.^80^ Second, less frequent emptying would have created fewer opportunities for environmental fecal contamination to occur and hygienic emptying may have reduced the quantity of fecal sludge that contaminated soils during emptying. In addition, intervention systems contained a drain for bathing, which may have prevented fecally contaminated graywater from flowing into nearby soils, and the concrete floors were likely easier to clean than control systems with dirt floors.^56^

Although our findings suggest that some pathogens appeared to be reduced by the latrine improvements, it is likely that the potential for exposure remains high in this setting.^13^ While we detected some individual pathogens, such as *Ascaris lumbricoides*, EAEC, *Shigella*/EIECand EPEC, in intervention soils less frequently compared to controls during the 24-month phase, we also detected one or more enteric pathogens in 86% of intervention latrine entrance soils two years post-intervention. Fecal waste from children unable to use the latrines was not addressed by the intervention.^28,84^ At the 24-month follow-up, 29% (289/980) of children reported defecating into a latrine, 29% (281/980) defecated into a child potty which was emptied into a latrine, 20% (192/980) used disposable diapers that were disposed with solid waste, 7.3% defecated on the ground (72/980), and 2.7% (26/980) defecated into diapers that were washed and reused (Table S6). In addition, the intervention did not address animal feces. While we adjusted for animals in our DID estimates, many animals are not penned in this setting and may defecate outside of their respective compounds, which was not accounted for in our analysis.^30^

Live chickens are also commonly purchased and stored in the compound for consumption.^85^ We may not have adequately captured this intermittent chicken ownership in our cross-sectional surveys.

The similar reduction in pathogen detection in soils and child stools may be informative about exposures. At two years post-intervention in the MapSan cohort, children born into study compounds were 1-24 months old, while children born previously and enrolled at baseline were 25-73 months old.^58^ Considering the consistent reduction in the detection of pathogens observed in soils and stools from children 1-24 months old, the dominant exposure pathways for these younger children may be inside the compound or soil ingestion may have represented a more important transmission pathway for these children.^86^ Older children are more mobile than younger children, and their potential exposures outside of study compounds may explain why the intervention did not reduce the prevalence of pathogen carriage among them.

Our study had several limitations, including a relatively small sample size that was not intended to observe small reductions in pathogen detection. Nevertheless, in high burden settings, sanitation interventions may need to achieve a large reduction in environmental fecal contamination both within households and in the larger community to reduce exposure risks and yield improved health outcomes.^87^ Further, intervention compounds had lower wealth indices and higher compound populations 24-months post intervention compared to control. This may suggest we underestimated changes due to sanitation improvements, but we adjusted for these in our regression analyses and did not observe substantial differences between unadjusted and adjusted point estimates that would indicate confounding. In addition, we assessed gene targets via molecular assays – which may not be 100% sensitive or specific^61,88,89^ – and not pathogen viability or infectivity.

There is substantial evidence that city-wide upgrades to sewerage infrastructure improve health outcomes.^74,75,90^ However, the high capital and maintenance costs^91^, and water usage requirements^92^ of such improvements suggest they are currently impractical for many LMICs. Until sewerage becomes widely feasible in high-burden settings, onsite sanitation systems remain necessary to achieve safely managed sanitation in many urban areas. The results of this study – and other rigorous environmental impact evaluations of onsite sanitation interventions^18,38,77^ – suggest that fecal contamination is transported into the environment through multiple complex pathways that may vary among settings.^93^ In urban Maputo – and similar settings with poor sanitation infrastructure, widespread environmental fecal contamination, and a high burden of enteric infection – other, more transformative interventions interrupting multiple transmission pathways may need to accompany improvements to onsite sanitation infrastructure. These improvements likely require an integrated and incremental approach that might include legal protections (e.g. land tenure)^94^, contact control interventions (e.g. hardscape cleanable flooring)^13,95,96^, public infrastructure (e.g. drainage, and improvements in quality, quantity, and access to water)^97^, and public services (e.g. education, hygienic fecal sludge and solid waste management)^20,98,99^. Such improvements may reduce the transport of enteric pathogens into the environment through site-specific pathways and subsequently reduce children’s infection risks.

## Supplemental Information

1. Text S1. Detailed description of the sanitation intervention

2. Text S2. Compound enrollment at baseline

3. Text S3. Test for Matrix Inhibition

4. Text S4. Custom TaqMan Array Card (TAC)

5. Table S1. Assays used on the custom TAC

6. Table S2. Interpretation of gene targets on the TAC

7. Table S3. Description of variables and their respective sources

8. Table S4. Soils samples matched at baseline and 24-month trial periods

9. Figure S1. Histogram of dates that latrine entrance soils were collected

10. Table S5. Mean Cq Values

11. Table S6. Child feces disposal at 24-month phase

## Notes

The findings and conclusions in this report are those of the authors and do not necessarily represent the official position of the Centers for Disease Control and Prevention.

## Notes

### Competing Interest Statement

The authors have declared no competing interest.

