## Supporting Information for "Impact of an Urban Sanitation Intervention on Enteric Pathogen Detection in Soils"

Supplemental Information

1. Text S1. Detailed description of the sanitation intervention
2. Text S2. Compound enrollment at baseline
3. Text S3. Test for Matrix Inhibition
4. Text S4. Custom TaqMan Array Card (TAC)
5. Table S1. Assays used on the custom TAC
6. Table S2. Interpretation of gene targets on the TAC
7. Table S3. Description of variables and their respective sources
8. Table S4. Soils samples matched at baseline and 24-month trial periods
9. Figure S1. Histogram of dates that latrine entrance soils were collected
10. Table S5. Mean Cq Values
11. Table S6. Child feces disposal at 24-month phase

Text S1. Detailed description of the sanitation intervention

WSUP initially built 250 shared toilets and 50 community sanitation blocks, but due to the depreciation of the Metical (Mozambican currency) in 2016-2017, WSUP built 150 additional shared toilets in the project area. Shared latrines became the property of the residents and included a toilet, superstructure, septic tank, and a lined infiltration pit. Community sanitation blocks officially remained the property of the municipality and included the same infrastructure as a shared latrine, but contained multiple toilets (one toilet per twenty people), a new piped water connection with a water storage tank, sink pedestal for handwashing (no running water but the drain was connected to the septic tank), rainwater harvesting tank, cement laundry basin, and community sanitation blocks with ≥60 residents received a urinal on an external wall of the structure which drained to the septic tank. Compound residents that received community sanitation blocks formed sanitation management committees, which were responsible for maintaining the sanitation infrastructure. The septic tanks in the shared latrines and community sanitation blocks were sized according to the number of users and were designed to be emptied every two years (assuming 40 liters accumulation per person per year). All intervention septic tanks contained an access port for hygienic emptying, but the ports were sealed shut and did not enable easy visual inspection of fecal sludge levels.

Text S2. Compound enrollment at baseline

At study baseline we intended to enroll an equal number of control and intervention compounds for soil sample collection. We mistakenly collected soil samples from more control compounds (n = 51) compared to compounds which received an intervention after the baseline visit (n=49) due to a miscommunication between the implementing organization, the research team, and the field staff.

Text S3. Test for Matrix Inhibition

We used the TaqMan™ Exogenous Internal Positive Control Assay to test soils samples for matrix inhibition. Following the manufacturer's protocol, we spiked in the internal positive control (IPC) to qPCR assays containing IPC specific primers and probes, TaqMan Universal PCR Mastermix, and nucleic acid extract from each soil sample. This assay is not designed for normalization, but it is designed to determine if a sample exhibits PCR inhibition. A negative call for the IPC suggests the presence of PCR inhibition and positive call for the IPC suggests an absence of PCR inhibition. We ran this assay on an Applied Biosystems 7500 real-time PCR system with manufacturer recommended thermocycling conditions: two minutes at 50°C, ten minutes at 95°C, followed by 40 cycles of 95°C for 15 seconds followed by 60°C for one minute. We observed positive amplification of the IPC in all assays suggesting an absence of PCR inhibition in our samples. The lack of PCR inhibition may be a result of the mass of soil we extracted from. The RNeasy PowerSoil Total RNA Kit and RNeasy PowerSoil DNA Elution Kit are designed for extraction from up to two grams of soil, but we only extracted from one gram of dry weight soil. This relatively low mass of soil used may have reduced the potential for inhibitors in our extracts.

Text S4. Custom TaqMan Array Card (TAC)

We purchased custom TACs produced by Thermo Fisher Scientific (Waltham, MA). TAC is a 384-well array card with 8 ports for loading samples and each 1-µL well contains dried-down primers and hydrolysis probes for the detection of defined targets.

For analysis, we mixed 25 µL of DNA template and 25 µL of RNA template (0.5 µL total template per reaction well) with 50 µL of qScript XLT 1-Step RT-qPCR ToughMix (Quantabio, Beverly, MA), then filled ports 2-7 with the combined 100 µL. In total we tested 6 samples per card, using the first port as a negative control and the last port as a positive control, for which we used individual aliquots of our combined positive control material (gene targets inserted into plasmids) (IDT, Coralville, IA). Combined positive controls were developed using methods from Kodani *et al.* 2012. Following the manufacturer's instructions, we centrifuged each card twice at 1,200 rpm for one minute, sealed the card, trimmed the loading ports, and loaded the card into a QuantStudio 7 (Thermo Fisher Scientific, Waltham, MA). All positive controls amplified as expected (typically ~ Cq = 28-30 depending on the assay) and we detected MS2 in all samples. Among 16 extraction controls and 16 no template controls we observed no amplification for any target below a Cq of 40.

Using the extraction methods described in the manuscript, the 0.25 µL of DNA or RNA template in each reaction well on TAC represents a 400-fold dilution from the starting gram of soil.

Equation S1: $dilution\ factor\ = \frac{0.25\ \mu\text{L template per well}}{100\ \mu\text{L total}} \times$

$1\ gram\ dry\ weight\ soil = \frac{1}{400}$

Table S1. Assays used on the custom TAC

| Target | Assay reference |
|---|---|
| ***Bacteria*** | |
| *Campylobacter coli* | Cunningham, S. A.; Sloan, L. M.; Nyre, L. M.; Vetter, E. A.; Mandrekar, J.; Patel, R. Three-Hour Molecular Detection of Campylobacter, Salmonella, Yersinia, and Shigella Species in Feces with Accuracy as High as That of Culture. *J. Clin. Microbiol.* **2010**, *48* (8), 2929–2933. |
| *Campylobacter jejuni* | Cunningham, S. A.; Sloan, L. M.; Nyre, L. M.; Vetter, E. A.; Mandrekar, J.; Patel, R. Three-Hour Molecular Detection of Campylobacter, Salmonella, Yersinia, and Shigella Species in Feces with Accuracy as High as That of Culture. *J. Clin. Microbiol.* **2010**, *48* (8), 2929–2933. |
| *Clostridium difficile* (*tcdA*) | Houser, B. A.; Hattel, A. L.; Jayarao, B. M. Real-Time Multiplex Polymerase Chain Reaction Assay for Rapid Detection of Clostridium Difficile Toxin-Encoding Strains. *Foodborne Pathog. Dis.* **2010**, *7* (6), 719–726. |
| *Clostridium difficile* (*tcdB*) | Houser, B. A.; Hattel, A. L.; Jayarao, B. M. Real-Time Multiplex Polymerase Chain Reaction Assay for Rapid Detection of Clostridium Difficile Toxin-Encoding Strains. *Foodborne Pathog. Dis.* **2010**, *7* (6), 719–726. |
| *E. coli / Shigella* (*ipaH* gene) | Thiem, V. D.; Sethabutr, O.; Seidlein, L. von; Tung, T. Van; Canh, D. G.; Chien, B. T.; Tho, L. H.; Lee, H.; Houng, H.-S.; Hale, T. L.; et al. Detection of Shigella by a PCR Assay Targeting the IpaH Gene Suggests Increased Prevalence of Shigellosis in Nha Trang, Vietnam. *J. Clin. Microbiol.* **2004**, *42* (5), 2031–2035. |
| EAEC (*aaiC* gene) | Liu, J.; Gratz, J.; Amour, C.; Kibiki, G.; Becker, S.; Janaki, L.; Verweij, J. J.; Taniuchi, M.; Sobuz, S. U.; Haque, R.; et al. A Laboratory-Developed Taqman Array Card for Simultaneous Detection of 19 Enteropathogens. *J. Clin. Microbiol.* **2013**, *51* (2), 472–480. |
| EAEC (*aatA* gene) | Boisen, N.; Struve, C.; Scheutz, F.; Krogfelt, K. A.; Nataro, J. P. New Adhesin of Enteroaggregative Escherichia Coli Related to the Afa/Dr/AAF Family. *Infect. Immun.* **2008**, *76* (7), 3281–3292. |
| EPEC (*bfpA* gene) | Liu, J.; Gratz, J.; Amour, C.; Kibiki, G.; Becker, S.; Janaki, L.; Verweij, J. J.; Taniuchi, M.; Sobuz, S. U.; Haque, R.; et al. A Laboratory-Developed Taqman Array Card for Simultaneous Detection of 19 Enteropathogens. *J. Clin. Microbiol.* **2013**, *51* (2), 472–480. |
| EPEC (*eae* gene) | Liu, J.; Gratz, J.; Amour, C.; Kibiki, G.; Becker, S.; Janaki, L.; Verweij, J. J.; Taniuchi, M.; Sobuz, S. U.; Haque, R.; et al. A Laboratory-Developed Taqman Array Card for Simultaneous Detection of 19 Enteropathogens. *J. Clin. Microbiol.* **2013**, *51* (2), 472–480. |
| ETEC-LT | Hidaka, A.; Hokyo, T.; Arikawa, K.; Fujihara, S.; Ogasawara, J.; Hase, A.; Hara-Kudo, Y.; Nishikawa, Y. Multiplex Real-Time PCR for Exhaustive Detection of Diarrhoeagenic *Escherichia Coli*. *J. Appl. Microbiol.* **2009**, *106* (2), 410–420. |
| ETEC-ST | Liu, J.; Gratz, J.; Amour, C.; Kibiki, G.; Becker, S.; Janaki, L.; Verweij, J. J.; Taniuchi, M.; Sobuz, S. U.; Haque, R.; et al. A Laboratory-Developed Taqman Array Card for Simultaneous Detection of 19 Enteropathogens. *J. Clin. Microbiol.* **2013**, *51* (2), 472–480. |

| | |
|---|---|
| *Salmonella* | Liu, J.; Gratz, J.; Amour, C.; Kibiki, G.; Becker, S.; Janaki, L.; Verweij, J. J.; Taniuchi, M.; Sobuz, S. U.; Haque, R.; et al. A Laboratory-Developed Taqman Array Card for Simultaneous Detection of 19 Enteropathogens. *J. Clin. Microbiol.* **2013**, *51* (2), 472–480. |
| Shiga-like toxin 1 (*stx1*) | Liu, J.; Gratz, J.; Amour, C.; Kibiki, G.; Becker, S.; Janaki, L.; Verweij, J. J.; Taniuchi, M.; Sobuz, S. U.; Haque, R.; et al. A Laboratory-Developed Taqman Array Card for Simultaneous Detection of 19 Enteropathogens. *J. Clin. Microbiol.* **2013**, *51* (2), 472–480. |
| Shiga-like toxin 2 (*stx2*) | Hidaka, A.; Hokyo, T.; Arikawa, K.; Fujihara, S.; Ogasawara, J.; Hase, A.; Hara-Kudo, Y.; Nishikawa, Y. Multiplex Real-Time PCR for Exhaustive Detection of Diarrhoeagenic *Escherichia Coli*. *J. Appl. Microbiol.* **2009**, *106* (2), 410–420. |
| *Vibrio cholerae* | Liu, J.; Gratz, J.; Amour, C.; Kibiki, G.; Becker, S.; Janaki, L.; Verweij, J. J.; Taniuchi, M.; Sobuz, S. U.; Haque, R.; et al. A Laboratory-Developed Taqman Array Card for Simultaneous Detection of 19 Enteropathogens. *J. Clin. Microbiol.* **2013**, *51* (2), 472–480. |
| *Yersinia* spp. | Liu, J.; Gratz, J.; Maro, A.; Kumburu, H.; Kibiki, G.; Taniuchi, M.; Howlader, A. M.; Sobuz, S. U.; Haque, R.; Talukder, K. A.; et al. Simultaneous Detection of Six Diarrhea-Causing Bacterial Pathogens with an In-House PCR-Luminex Assay. *J. Clin. Microbiol.* **2012**, *50* (1), 98–103. |
| **Viruses** | |
| Adenovirus 40/41 | Jothikumar, N.; Cromeans, T. L.; Hill, V. R.; Lu, X.; Sobsey, M. D.; Erdman, D. D. Quantitative Real-Time PCR Assays for Detection of Human Adenoviruses and Identification of Serotypes 40 and 41. *Appl. Environ. Microbiol.* **2005**, *71* (6), 3131–3136. |
| Astrovirus | Liu, J.; Kibiki, G.; Maro, V.; Maro, A.; Kumburu, H.; Swai, N.; Taniuchi, M.; Gratz, J.; Toney, D.; Kang, G.; et al. Multiplex Reverse Transcription PCR Luminex Assay for Detection and Quantitation of Viral Agents of Gastroenteritis. *J. Clin. Virol.* **2011**, *50* (4), 308–313. |
| Norovirus GI | Jothikumar, N.; Lowther, J. A.; Henshilwood, K.; Lees, D. N.; Hill, V. R.; Vinjé, J. Rapid and Sensitive Detection of Noroviruses by Using TaqMan-Based One-Step Reverse Transcription-PCR Assays and Application to Naturally Contaminated Shellfish Samples. *Appl. Environ. Microbiol.* **2005**, *71* (4), 1870–1875. |
| Norovirus GII | Kageyama, T.; Kojima, S.; Shinohara, M.; Uchida, K.; Fukushi, S.; Hoshino, F. B.; Takeda, N.; Katayama, K. Broadly Reactive and Highly Sensitive Assay for Norwalk-like Viruses Based on Real-Time Quantitative Reverse Transcription-PCR. *J. Clin. Microbiol.* **2003**, *41* (4), 1548–1557. |
| Rotavirus A | Jothikumar, N.; Kang, G.; Hill, V. R. Broadly Reactive TaqMan® Assay for Real-Time RT-PCR Detection of Rotavirus in Clinical and Environmental Samples. *J. Virol. Methods* **2009**, *155* (2), 126–131. |
| Sapovirus I/II/IV | Liu, J.; Gratz, J.; Amour, C.; Kibiki, G.; Becker, S.; Janaki, L.; Verweij, J. J.; Taniuchi, M.; Sobuz, S. U.; Haque, R.; et al. A Laboratory-Developed Taqman Array Card for Simultaneous Detection of 19 Enteropathogens. *J. Clin. Microbiol.* **2013**, *51* (2), 472–480. |

| | |
|---|---|
| Sapovirus V | Liu, J.; Gratz, J.; Amour, C.; Kibiki, G.; Becker, S.; Janaki, L.; Verweij, J. J.; Taniuchi, M.; Sobuz, S. U.; Haque, R.; et al. A Laboratory-Developed Taqman Array Card for Simultaneous Detection of 19 Enteropathogens. *J. Clin. Microbiol.* **2013**, *51* (2), 472–480. |
| ***Protozoa*** | |
| *Cryptosporidium parvum* | Jothikumar, N.; da Silva, A. J.; Moura, I.; Qvarnstrom, Y.; Hill, V. R. Detection and Differentiation of Cryptosporidium Hominis and Cryptosporidium Parvum by Dual TaqMan Assays. *J. Med. Microbiol.* **2008**, *57* (9), 1099–1105. |
| *Entamoeba histolytica* | Verweij, J. J.; Blangé, R. A.; Templeton, K.; Schinkel, J.; Brienen, E. A. T.; van Rooyen, M. A. A.; van Lieshout, L.; Polderman, A. M. Simultaneous Detection of Entamoeba Histolytica, Giardia Lamblia, and Cryptosporidium Parvum in Fecal Samples by Using Multiplex Real-Time PCR. *J. Clin. Microbiol.* **2004**, *42* (3), 1220–1223. |
| *Giardia duodenalis* | Verweij, J. J.; Blangé, R. A.; Templeton, K.; Schinkel, J.; Brienen, E. A. T.; van Rooyen, M. A. A.; van Lieshout, L.; Polderman, A. M. Simultaneous Detection of Entamoeba Histolytica, Giardia Lamblia, and Cryptosporidium Parvum in Fecal Samples by Using Multiplex Real-Time PCR. *J. Clin. Microbiol.* **2004**, *42* (3), 1220–1223. |
| ***Soil-transmitted helminths*** | |
| *Ascaris lumbricoides* | Wiria, A. E.; Prasetyani, M. A.; Hamid, F.; Wammes, L. J.; Lell, B.; Ariawan, I.; Uh, H. W.; Wibowo, H.; Djuardi, Y.; Wahyuni, S.; et al. Does Treatment of Intestinal Helminth Infections Influence Malaria? Background and Methodology of a Longitudinal Study of Clinical, Parasitological and Immunological Parameters in Nangapanda, Flores, Indonesia (ImmunoSPIN Study). *BMC Infect. Dis.* **2010**, *10* (1), 77. |
| *Trichuris trichiuria* | Pilotte, N.; Papaiakovou, M.; Grant, J. R.; Bierwert, L. A.; Llewellyn, S.; McCarthy, J. S.; Williams, S. A. Improved PCR-Based Detection of Soil Transmitted Helminth Infections Using a Next-Generation Sequencing Approach to Assay Design. *PLoS Negl. Trop. Dis.* **2016**, *10* (3), e0004578. |

Table S2. Interpretation of gene targets on the TAC

| Target | Gene Targeted | Interpretation |
|---|---|---|
| **Bacteria** | | |
| *Campylobacter coli* | *cadF* gene | If either was detected, call as *Campylobacter coli/jejuni* positive |
| *Campylobacter jejuni* | *cadF gene* | |
| *Clostridium difficile* (*tcdA*) | *tcdA* gene | If either was detected, call as *Clostridium difficile* positive |
| *Clostridium difficile* (*tcdB)* | *tcdB gene* | |
| *E. coli / Shigella* (*ipaH*) | *ipaH* gene | If detected, call as *Shigella*/EIEC positive |
| EAEC (*aaiC*) | *aaiC* gene | If either was detected, call as EAEC positive |
| EAEC (*aatA*) | *aatA* gene | |
| EPEC (*bfpA*) | *bfpA* gene | If either was detected, call as EPEC positive |
| EPEC (*eae*) | *eae* gene | |
| ETEC-LT | *LT* gene | If either was detected, call as ETEC positive |
| ETEC-ST | *STh/STp* | |
| *Salmonella* spp. | *invA* gene | If detected, call as *Salmonella spp.* positive |
| Shiga-like toxin 1 (stx1) | *$stx_1$* gene | If either was detected, call as STEC positive |
| Shiga-like toxin 2 (stx2) | *$stx_2$* gene | |
| *Vibrio cholerae* | *toxR* gene | If detected, call as *Vibrio cholerae* positive |
| *Yersinia* spp. | *lysP* gene | If detected, call as *Yersinia spp.* positive |
| **Viruses** | | |
| Adenovirus 40/41 | *Fiber* gene | If detected, call as Adenovirus 40/41 positive |
| Astrovirus | *Capsid* gene | If detected, call as Astrovirus positive |
| Norovirus GI | *ORF1-ORF2* gene | If either was detected, call as Norovirus GI/GII positive |
| Norovirus GII | *ORF1-ORF2* gene | |
| Rotavirus A | *NSP3* gene | If detected, call as Rotavirus A positive |
| Sapovirus I/II/IV | *RdRp* gene | If either was detected, call as Sapovirus positive |
| Sapovirus V | *RdRp* gene | |
| ***Protozoa*** | | |
| *Cryptosporidium parvum* | *18S* | If detected, call as *Cryptosporidium parvum* positive |
| *Entamoeba histolytica* | *18S* | If detected, call as *Entamoeba histolytica* positive |
| *Giardia duodenalis* | *18S* | If detected, call as *Giardia duodenalis* positive |

| ***Helminth*** | | |
|---|---|---|
| *Ascaris lumbricoides* | *18S* | If detected, call as *Ascaris lumbricoides* positive |
| *Trichuris trichiuria* | *ITS1* | If detected, call as *Trichuris trichiuria* positive |

Table S3. Description of variables and their respective sources

| | **Variable description** | **Data source** |
|---|---|---|
| **Outcome Data** | | |
| Presence of ≥1 enteric pathogen in latrine entrance soils | Binary detect/non-detect; 1/0 | Experimental data |
| Total number of enteric pathogens detected | Count; from 0 to 20 | Experimental data |
| Presence of individual pathogens on TAC | Binary detect/non-detect; 1/0 | Experimental data |
| **Covariates used in multivariate model selection** | | |
| Compound population | Continuous variable: transformed to represent a 10-person increase | Baseline and 24-month datasets |
| Wealth index | Quartile (1, 2, 3, or 4) derived from a continuous variable (from 0 to 1) | Baseline and 24-month datasets (Calculated using the Simple Poverty Scorecard® Poverty-Assessment Tool: Mozambique) |
| Visibly wet soil | Wet/dry; 1/0 | Observed and recorded by enumerator at time of sampling |
| Sun exposure status | Factor; complete sun, partially shaded, complete shade | Observed and recorded by enumerator at time of sampling |
| Average temperature in Fahrenheit during the day of and day before the soil sample was collected (e.g. 2-day average temperature) | Continuous variable, mean centered | Downloaded data from the National Oceanic and Atmospheric Administration's National Centers for Environmental Information |

| | | |
|---|---|---|
| | | (https://www.ncdc.noaa.gov/cdo-web/datatools/findstation) |
| Baseline and 24-month sanitation infrastructure | Factor; Pit latrine (without slab), pit latrine (with slab), intervention pour-flush toilet, non-intervention pour flush toilet, or unusable latrine (e.g. used neighbor's latrine or reported open defecation) | Baseline and 24-month datasets<br><br>In addition, we reviewed illustrative photographs of sanitation infrastructure to confirm the sanitation infrastructure present |
| Dog(s) present | Binary, present / not present; 1/0 | Baseline and 24-month datasets |
| Chicken(s)/duck(s) present | Binary, present / not present; 1/0 | Baseline and 24-month datasets |
| Cat(s) present | Binary, present / not present; 1/0 | Baseline and 24-month datasets |
| Visible feces in the compound (human or animal) | Binary, present / not present; 1/0 | Baseline and 24-month datasets |

Table S4. Soils samples matched at baseline and 24-month trial periods

| **Latrine entrance soil samples** | **Control** | **Intervention** |
|---|---|---|
| Just baseline | 6 | 2 |
| Matched baseline and 24-month | 42 | 41 |
| Just 24-month | 3 | 2 |
| Total baseline | 48 | 45 |
| Total 24-month | 43 | 43 |

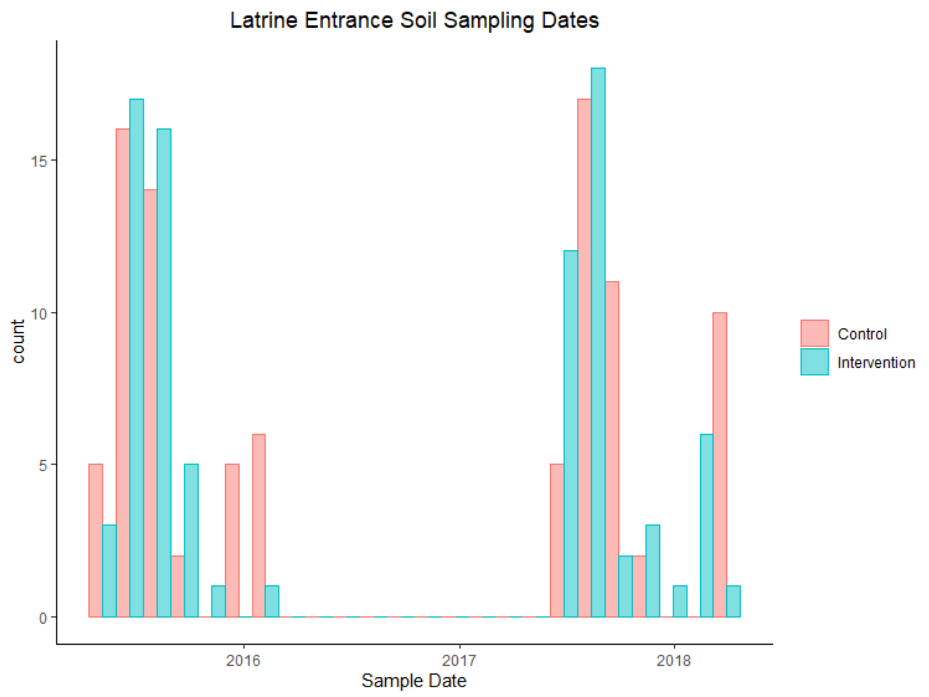

Figure S1. Histogram of dates that latrine entrance soils were collected

Table S5. Mean Cq Values

| **Pathogen** | Baseline Cq | 24-month Cq |
|---|---|---|
| *Ascaris lumbricoides* | | |
| control | 28.4 (4.6) | 29.6 (4.1) |
| intervention | 29.0 (4.4) | 28.3 (4.2) |
| Enteroaggregative *E. coli* | | |
| control | 31.9 (2.0) | 31.0 (2.8) |
| intervention | 32.7 (1.8) | 32.2 (2.6) |
| *Giardia duodenalis* | | |
| control | 30.9 (2.7) | 32.5 (2.7) |
| intervention | 32.9 (2.9) | 33.1 (2.7) |
| *Shigella*/Enteroinvasive *E. coli* | | |
| control | 32.6 (1.3) | 32.4 (2.5) |
| intervention | 34.9 (2.5) | 33.3 (0.97) |
| Enterotoxigenic *E. coli* | | |
| control | 33.8 (4.0) | 33.7 (3.1) |
| intervention | 34.4 (3.4) | 33.4 (2.5) |
| adenovirus 40/41 | | |
| control | 30.9 (2.8) | 31.6 (2.2) |
| intervention | 30.9 (4.6) | 31.0 (1.7) |
| astrovirus | | |
| control | 32.8 (4.4) | 33.0 (3.2) |
| intervention | 34.3 (4.2) | 33.7 (3.7) |
| Enteropathogenic *E. coli* | | |
| control | 31.5 (2.3) | 32.0 (2.33) |
| intervention | 31.0 (3.8) | 31.2 (2.4) |
| *Trichuris trichiura* | | |
| control | 27.6 (4.3) | 26.8 (4.0) |
| intervention | 28.5 (3.7) | 28.5 (5.6) |
| *Clostridium difficile* | | |
| control | 34.3 (0.87) | 33.8 (1.4) |
| intervention | 32.7 (2.8) | 34.2 (1.0) |
| *Cryptosporidium parvum* | | |
| control | NA | |
| intervention | | |
| *Salmonella* spp. | | |
| control | NA | |
| intervention | | |
| Shiga-toxin producing *E. coli* | | |
| control | NA | |
| intervention | | |

| | |
|---|---|
| *Campylobacter jejuni/coli* | |
| control | NA |
| intervention | |
| *Yersinia* spp. | |
| control | NA |
| intervention | |
| norovirus GI/GII | |
| control | NA |
| intervention | |
| rotavirus A | |
| control | NA |
| intervention | |
| *Entamoeba histolytica* | |
| control | NA |
| intervention | |
| sapovirus I/II/IV/V | |
| control | NA |
| intervention | |
| *Vibrio cholerae* | |
| control | NA |
| intervention | |

Note: Cq values are the mean of detected samples and non-detects were not included in the calculation

Table S6. Child feces disposal at 24-month phase

| Feces Disposal | Sub-category | Survey response | |
|---|---|---|---|
| In a diaper | | 22% (218/980) | |
| | Diaper is washed and reused | | 2.7% (26/980) |
| | Diaper is discarded with solid waste | | 20% (192/980) |
| In the latrine | | 29% (289/980) | |
| | In the latrine | | 29% (289/980) |
| On the ground | | 7.3% (72/980) | |
| | Left on the ground | | 0.3% (3/980) |
| | Put with the solid waste | | 0.1% (1/980) |
| | Put into a soakaway pit | | 0.1% (1/980) |
| | Put into the latrine | | 5.6% (55/980) |
| | Buried | | 1.2% (12/980) |
| Child potty (contents emptied into the latrine) | | 29% (281/980) | |
| | Child potty | | 29% (281/980) |
| No response | | 12% (120/980) | |
| | No response | | 12% (120/980) |
